# Alveolar macrophages up-regulate a non-classical innate response to *Mycobacterium tuberculosis* infection *in vivo*

**DOI:** 10.1101/520791

**Authors:** A. C. Rothchild, G. S. Olson, J. Nemeth, L. M. Amon, D. Mai, E. S. Gold, A. H. Diercks, A. Aderem

**Affiliations:** Seattle Children’s Research Institute, Center for Global Infectious Disease Research, Seattle, WA 98109.; Medical Scientist Training Program, University of Washington School of Medicine, Seattle, WA 98195.; University Hospital Zurich, University of Zurich, Division of Infectious Diseases and Hospital Epidemiology, Raemistrasse 100, 8091 Zürich, Switzerland; Universal Cells, Seattle, WA 98121.

## Abstract

Alveolar macrophages (AMs) are the first cells to be infected during *Mycobacterium tuberculosis* (Mtb) infection. Thus the AM response to infection is the first of many steps leading to initiation of the adaptive immune response, which is required for efficient control of infection. A hallmark of Mtb infection is the delay of the adaptive response, yet the mechanisms responsible for this delay are largely unknown. We developed a system to identify, sort and analyze Mtb-infected AMs from the lung within the first 10 days of infection. In contrast to what has been previously described using *in vitro* systems, we find that Mtb-infected AMs up-regulate a cell-protective antioxidant transcriptional signature that is dependent on the lung environment and not dependent on bacterial virulence. Computational approaches including pathway analysis and transcription factor binding motif enrichment analysis identify Nrf2 as a master regulator of the response of AMs to Mtb infection. Using knock-out mouse models, we demonstrate that Nrf2 drives the expression of the cell protective transcriptional program and impairs the ability of the host to control bacterial growth over the first 10 days of infection. Mtb-infected AMs exhibit a highly delayed pro-inflammatory response, and comparisons with uninfected AMs from the same infected animals demonstrate that inflammatory signals in the lung environment are blocked in the Mtb-infected cells. Thus, we have identified a novel lung-specific transcriptional response to Mtb infection that impedes AMs from responding rapidly to intracellular infection and thereby hinders the overall immune response.

**One Sentence Summary:** In response to Mtb infection *in vivo*, alveolar macrophages fail to up-regulate the canonical pro-inflammatory innate response and instead induce an Nrf2-dependent cell protective transcriptional program, which in turn impairs the host’s control of bacterial growth.

## Introduction

Transmission of *Mycobacterium tuberculosis* (Mtb), the deadliest infectious pathogen worldwide, generally occurs via aerosols expelled by cough from an infected person. Although inhaled Mtb is rapidly engulfed by alveolar macrophages (AMs) in the lung (*1*), mobilization of a robust immune response is significantly delayed, which is thought to contribute to disease progression. In animal models, recruitment of innate cells such as neutrophils and monocytes to the lung and T cell priming in the draining mediastinal lymph nodes does not occur until 11-14 days following infection, by which time the bacteria have replicated nearly 1,000-fold in the lung (*1–5*). The mechanisms that restrain or subvert the host response in the lung immediately following infection remain largely unknown.

AMs are the first cells in the lung to be infected with Mtb after aerosol transmission (*1*). Originating from fetal monocytes, AMs are the largest resident macrophage population in the lung and are situated within individual alveoli between the airway and underlying Type I alveolar epithelial cells (*6*). AMs serve as pulmonary immune sentinels, constantly sampling the airway for foreign particles. In addition, AMs perform a critical homeostatic role clearing inhaled material from the airway and recycling pulmonary surfactant. Therefore, AMs must be able to uncouple phagocytic functions from inflammatory responses (*7, 8*). In the absence of functional AMs, both mice and humans suffer from a form of pulmonary inflammation known as pulmonary alveolar proteinosis (PAP), caused by the buildup of pulmonary surfactant (*9*). Due to these steady-state functions, AMs express unique transcriptional and epigenetic profiles that are highly distinct from those of other tissue-resident macrophages (*10–12*).

Similar to other macrophage populations that are constantly exposed to environmental stimuli, AMs express a number of inhibitory receptors that have been shown to dampen their responses. However, under conditions such as acute lung injury, AMs can become highly activated and release damaging levels of pro-inflammatory mediators that contribute to airway disease (*6*). During other pulmonary infections such as influenza, in which AMs serve as secondary or tertiary responders, AMs exhibit pro-inflammatory transcriptional responses and can serve as an immunotherapy target to limit inflammation (*13, 14*). However, the AM response is likely influenced by cytokines released into the pulmonary environment by other cell types during these infections.

Unlike most pulmonary infections, the initial infectious dose of Mtb is very low and for the first several days AMs are the only cells infected with the bacteria. Therefore, Mtb infection provides an opportunity to assess the ability of AMs to respond to direct intracellular infection rather than environmental cues. Results from two recent studies provide hints that the response of AMs to Mtb might be suboptimal: At two weeks following infection, when Mtb can be found in multiple cell populations, AMs were significantly less effective at controlling bacterial replication than interstitial macrophages, another macrophage subset found in the lung (*15*); and AMs facilitated relocalization of the bacteria from the airway into the lung interstitium (*1*).

Due to the paucity of Mtb-infected AMs during the first few days following aerosol challenge, their presumed immediate response to the bacteria has been extrapolated from *in vitro* culture systems using models such as bone marrow derived macrophages (BMDMs). These studies have identified numerous sensing pathways and cytokine responses including TNF, Type I IFN, and IL-1β (*16–21*) that have been shown to play crucial roles in host immunity in both animal and human studies (*22*). However, experiments using knock-out mice have demonstrated that these mediators do not affect the course of disease during the first week of infection *in vivo* (*23–25*), despite the fact that they are up-regulated within hours by Mtb-infected macrophages *in vitro*.

In order to more precisely define the response of AMs to Mtb *in vivo*, we developed an infection model and isolation procedure that yields sufficient numbers of infected AMs to perform global systems-level analyses at any point within the first 10 days following infection. Our results demonstrate that the up-regulation of genes normally associated with the macrophage response to Mtb *in vitro* is highly delayed in AMs infected *in vivo*. Immediately following *in vivo* infection, AMs up-regulate a cell-protective antioxidant transcriptional program regulated by the transcription factor, Nrf2. Activation of this program is dependent on the lung microenvironment and shapes the course of disease. In the absence of Nrf2, AMs have an enhanced ability to control bacterial growth through the first 10 days of infection.

## Results

### Alveolar macrophages are the first cells infected by Mtb in the lung

In a standard low-dose Mtb aerosol challenge model, mice are infected with ∼100 CFU, which, due to the low replication rate of the bacteria, does not allow for the isolation of sufficient numbers of Mtb-infected cells from a reasonable number of animals for analysis. Therefore, we developed a high-dose model in which mice are infected with 2 - 4 x 10^3^ CFU of an mEmerald-expressing H37Rv strain of Mtb. We analyzed the population of infected cells in the lung over the first two weeks using this model and found that the vast majority of infected cells early on were AMs with significant numbers of infected neutrophils (PMN) and monocytes-derived macrophages (MDMs) appearing between day 10 and 14 (**Fig 1A, S1**). The localization of the Mtb and the timing of recruitment of innate immune cells in the high-dose model is consistent with other studies that have used a low-dose model (*1, 3*), suggesting that the high-dose infection does not alter the initial immune response in the lung. At this dose, ∼0.5-2% of AMs were infected (**Fig 1B**) and the total number of Mtb-infected AMs did not change significantly over the first 2 weeks. By day 14, the numbers of infected neutrophils and AMs were equivalent (**Fig 1C**).

**Figure 1:**
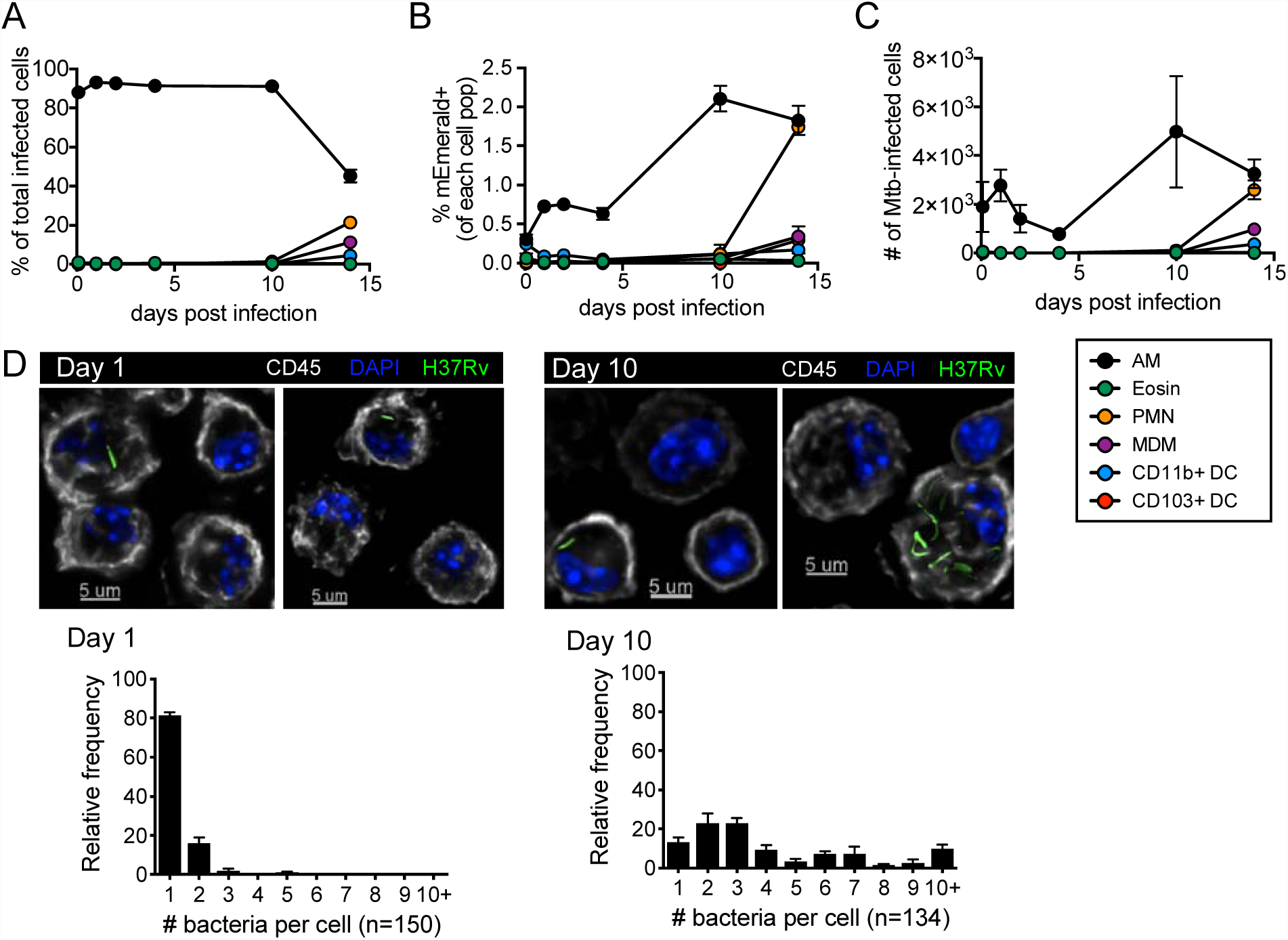
Alveolar macrophages are the first cells infected by Mtb after aerosol infection. (A) % of total infected cells, (B) % mEmerald^+^ for each cell population, and (C) total number of Mtb-infected cells in the lung between 2 hours and 14 days after high dose aerosol infection with mEmerald-tagged H37Rv (n = 3 mice/time point). (D) Microscopy of BAL samples 1 and 10 days after high dose aerosol infection with mEmerald-tagged H37Rv and quantitation of bacteria per AM (n = 3 replicates/time point, each replicate was pooled from 3 mice). *Abbreviations: AM = alveolar macrophages, PMN = neutrophils, Eosin = eosinophils, MDM = myeloid-derived macrophages, DC = dendritic cells.* Data are presented as mean ± SEM. Data is representative of 2 independent experiments.

We found that Mtb-infected AMs were enriched within bronchoalveolar lavage (BAL) samples compared to the total lung and used high-throughput microscopy to analyze BAL samples from mice following high-dose infection with mEmerald-H37Rv to quantify the number of bacteria in each infected cell. At 1 day post-infection, 81.3 ± 1.8% (mean ± SEM) of infected AMs contained a single bacillus; by day 10 only 13.2 ± 2.3% (mean ± SEM) of AMs contained a single bacillus, with 58.4 ± 6.1% of AMs containing 2-5 bacilli and 28.4 ± 3.8% containing more than 5 bacilli (**Fig 1D**). These data demonstrate that AM serve as a replication niche for Mtb through the first 10 days infection, an observation that has been similarly described at 1 and 2 weeks following infection in other studies (*1, 15*).

### Mtb-infected alveolar macrophages up-regulate an Nrf2-associated antioxidant response *in vivo*

To characterize the early macrophage transcriptional response to Mtb infection, we infected mice with 2 - 4 x 10^3^ CFU of mEmerald-H37Rv, and isolated AMs (CD45^+^, Zombie Violet^-^, CD3^-^/CD19^-^, Siglec-F^+^, CD11b^mid^, and CD64^+^) (*26, 27*) from BAL by fluorescence activated cell sorting (FACS) 24 hours following infection **(Fig S1)**. Isolating AMs by BAL allowed the samples to be kept on ice throughout processing and sorting, eliminating the need for lengthy and harsh digestion of lung tissue. Three populations were sorted for analysis by RNA-seq: Mtb-infected AMs (mEmerald^+^), bystander AMs (mEmerald^-^), and AMs from naïve mice **(Fig 2A).**

**Figure 2:**
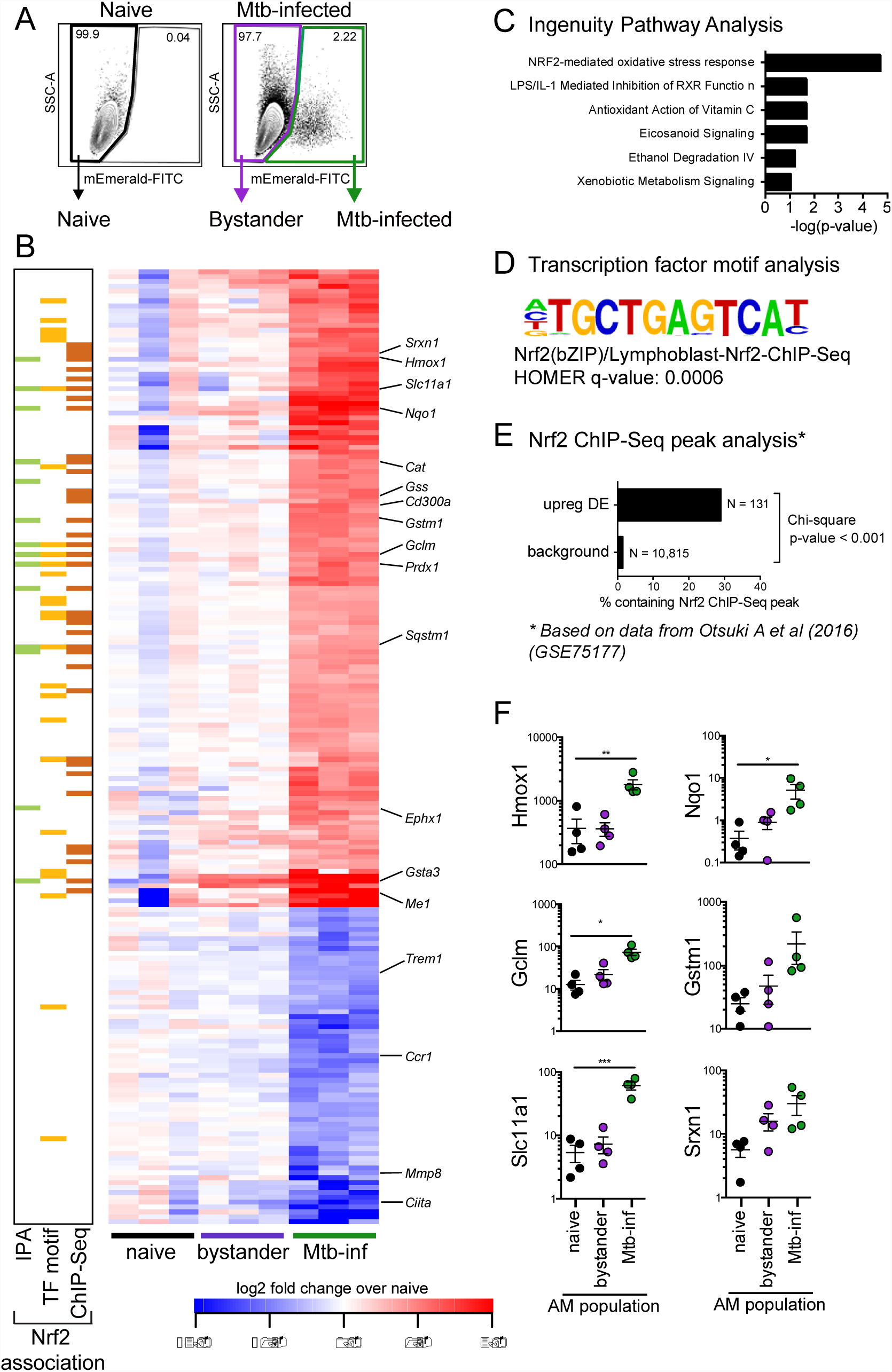
Mtb-infected alveolar macrophages up-regulate an Nrf2-associated antioxidant gene signature. (A) Gating scheme to sort naïve, bystander, and Mtb-infected AMs from bronchoalveolar lavage (BAL) samples after high dose aerosol infection with mEmerald-tagged H37Rv. (B) Heatmap of gene expression (log_2_ fold change over average of naïve AMs) for 196 differentially expressed genes between naïve and Mtb-infected AMs (Filtering criteria: average CPM >1, |fold change| > 2 and FDR < 0.01, Benjamini-Hochberg calculated). Columns are independent experiments (pooled mice) and rows are genes. Genes called out are known Nrf2 target genes of interest as well as downregulated pro-inflammatory genes. Colored bars to the left indicate Nrf2 association as determined by 3 different methods: (C) Ingenuity Pathway Analysis, (D) transcription factor promoter binding motif enrichment analysis (HOMER), and (E) ChIP-seq peak analysis. (F) RT-qPCR validation of Nrf2 associated genes for naïve, bystander and Mtb-infected AMs 24 hours post-infection. Values are relative to Ef1a. Data is presented from 3 independent experiments. Data are presented as mean ± SEM with one-way ANOVA with Dunnett’s post-test *p < 0.05, **p < 0.01, ***p < 0.001.

At 24 hours post-infection, we identified 196 genes that were differentially expressed (DE) (average counts per million (CPM) > 1, |fold change| > 2, FDR < 0.01) between Mtb-infected AMs and naïve AMs (**Fig 2B**). No genes were differentially expressed between bystander AMs and naïve AMs at this stringency, indicating an absence of systemic changes to the lung 24 hours after infection.

To identify enriched pathways and potential transcriptional regulators, we analyzed the up-regulated (131) and down-regulated (65) genes independently using three complementary computational approaches. Ingenuity Pathway Analysis (IPA) revealed that the up-regulated genes were significantly enriched for the “Nrf2-mediated oxidative stress response” pathway (p-value = 10^-11.3^) (**Fig 2B** (*green squares*), **Fig 2C**). This pathway includes genes involved in a number of cell protective functions including: antioxidant production (*Nqo1, Cat, Prdx1, Txnrd1*), iron metabolism (*Hmox1*), and glutathione metabolism (*Gstm1, Gclm, Gsta3*) (*28*). We also searched for enriched transcription factor promoter binding motifs using HOMER (Hypergeometric Optimization of Motif EnRichment) (*29*). The top four enriched transcription factor binding motifs in the 131 up-regulated genes (q-value = 0.0006) were motifs for either Nrf2 or Bach1, which are known to compete for the same binding sites (*30*) (**Fig 2B** (*yellow squares*), **Fig 2D)**. To validate these results, we used a published Nrf2 ChIP-Seq analysis of peritoneal macrophages that were stimulated with the Nrf2 agonist diethyl maleate (GSE75177) (*31*). 29% of the 131 up-regulated genes contained an Nrf2 ChIP-Seq peak, compared to 1.5% of background genes. The difference in proportions is highly significant, χ^2^ (1, N = 10,946) = 531.21, p-value < 2.2e^-16^) (**Fig 2B**, (*orange squares),* **Fig 2E**). We validated the up-regulation of several of the Nrf2-associated genes by qPCR (**Fig 2F**). These results demonstrate that the transcription factor Nrf2 is associated with the up-regulated transcriptional signature in Mtb-infected AMs 24 hours after infection *in vivo*. A similar analysis of the down-regulated genes did not uncover any significantly enriched pathways or candidate transcriptional regulators.

### Mtb-infected alveolar macrophages do not up-regulate classical pro-inflammatory genes

We also examined whether, in addition to Nrf2-associated genes, Mtb-infected AMs up-regulated classical pro-inflammatory genes that have been seen previously expressed by various macrophage subsets in response to Mtb including cytokines and chemokines (e.g. *Tnf, Il1b, Cxcl9)* (*22*), pro-inflammatory receptors, costimulatory molecules, and Fc receptors. Very few of these genes were significantly up-regulated by Mtb-infected AMs (**Fig. S2**). In contrast, several genes involved in glycolytic metabolism (*Hif1a, Pkm, Aldoa*) were significantly up-regulated by Mtb-infected AMs (*32*).

### Virulent Mtb is not required to activate the Nrf2-associated signature

To determine whether the Nrf2-signature was specifically induced by virulent Mtb, we repeated the high-dose infections using an mEmerald-expressing strain of Mtb lacking the RD1 virulence locus (ΔRD1-H37Rv) and using fluorescent 1μM carboxylate-coated latex beads (**Fig S3A, B)**. Both the ΔRD1-infected and bead-positive AMs displayed up-regulation of Nrf2-associated genes at 24 hours, although the magnitude of the fold change for some genes was smaller than for H37Rv-infected AMs (**Fig 3A)**. Unlike *in vitro* models that have shown a role for the ESX-1 locus, contained within RD1, in regulating the type I IFN responses of macrophages to Mtb (*16, 33, 34*), there were very few significant differences between the transcriptional responses of H37Rv vs ΔRD1-infected AMs 24 hours after infection (**Fig 3B).** The Nrf2-mediated oxidative stress response was the most highly enriched pathway in ΔRD1-infected AMs by IPA (**Table S1**). Based on the stringencies imposed on the 24 hour H37Rv-infected AMs, no genes were significantly changed in the bead-positive AMs and similarly no pathways were enriched by IPA (**Fig 3C, Table S1**). However, it is worth noting that within the most changed genes (with a filtering criteria of FDR < 0.05, fold change > 1.5) Nrf2-associated genes showed the greatest enrichment. Overall, these data demonstrate that AMs up-regulate the Nrf2-associated signature as a response to virulent bacteria, avirulent bacteria, or inert beads, suggesting that it is a more general response by AMs to the uptake of particles. While it is not surprising that particle uptake leads to transcriptional changes, it is notable that infection with a virulent pathogen appears to induce no additional host response within the first day of infection, suggesting that classical pathogen sensing is deficient, inhibited, or delayed in AMs.

**Figure 3:**
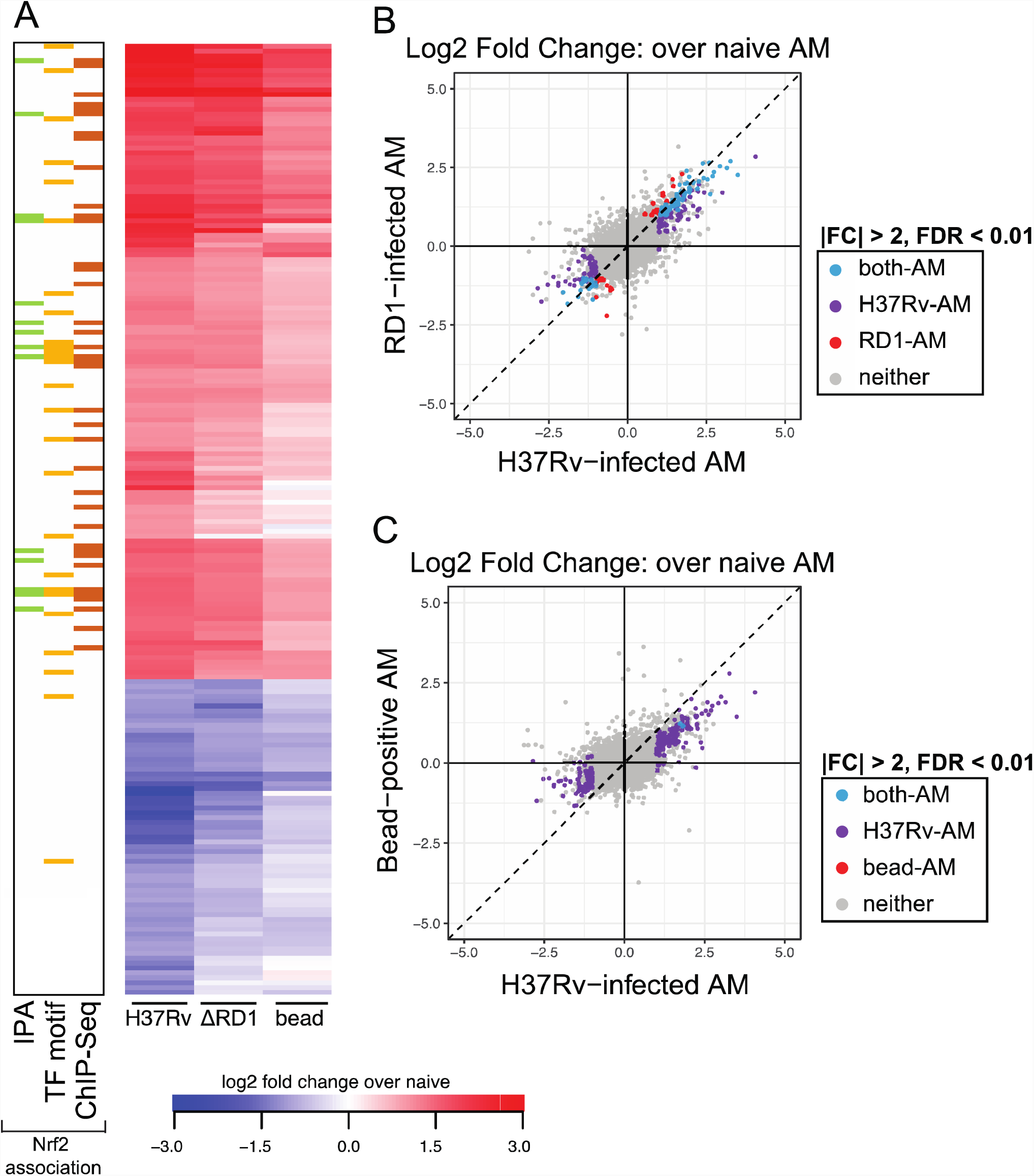
Up-regulation of Nrf2-associated signature does not require virulent Mtb infection. (A) Heatmap of log_2_ fold change gene expression over average of naïve AMs for H37Rv-infected, ΔRD1-infected, and bead-positive AMs 24 hours after treatment. Columns represent averages of 3 independent experiments. Rows represent 196 DE genes described in *Figure 2*. Colored bars to the left indicate Nrf2 association as described in *Figure 2*. (B, C) Scatterplots comparing gene expression values (log2 fold change over average of naïve AMs) for H37Rv-infected versus ΔRD1-infected AMs (B) or H37Rv-infected versus bead-positive AMs (C) 24 hours post-infection with significant differentially expressed genes highlighted (average CPM >1, |fold change| > 2 and FDR < 0.01, Benjamini-Hochberg calculated). Data is presented from 3 independent experiments.

### Over the first 10 days of infection, expression of Nrf2-associated genes is sustained and expression of pro-inflammatory genes is delayed in Mtb-infected alveolar macrophages

To characterize the kinetics of the AM response to Mtb infection, we extended our transcriptional analysis to include 4 additional time points: 0.5 (12 hours), 2, 4, and 10 days post-infection. We identified 288 genes that were significantly up-regulated at one or more of these time points compared to naïve AMs (**Fig 4A**). Many of the 131 genes up-regulated at 24 hours and associated with Nrf2 by either IPA, transcription factor motif analysis or ChIP-Seq analysis showed sustained expression through 10 days of infection (**Fig 4A,** *top*). Furthermore, by IPA, the “Nrf2-mediated oxidative stress response” pathway was the most highly enriched pathway at all 5 timepoints (p-values: 10^-5.3^-10^-10.1^) (**Table S1**). HOMER analysis also pinpointed a Nrf2 motif as the most enriched transcription factor motif at 2 4, and 10 days, and identified no enriched motif at 0.5 days **(Table S1).**

**Figure 4:**
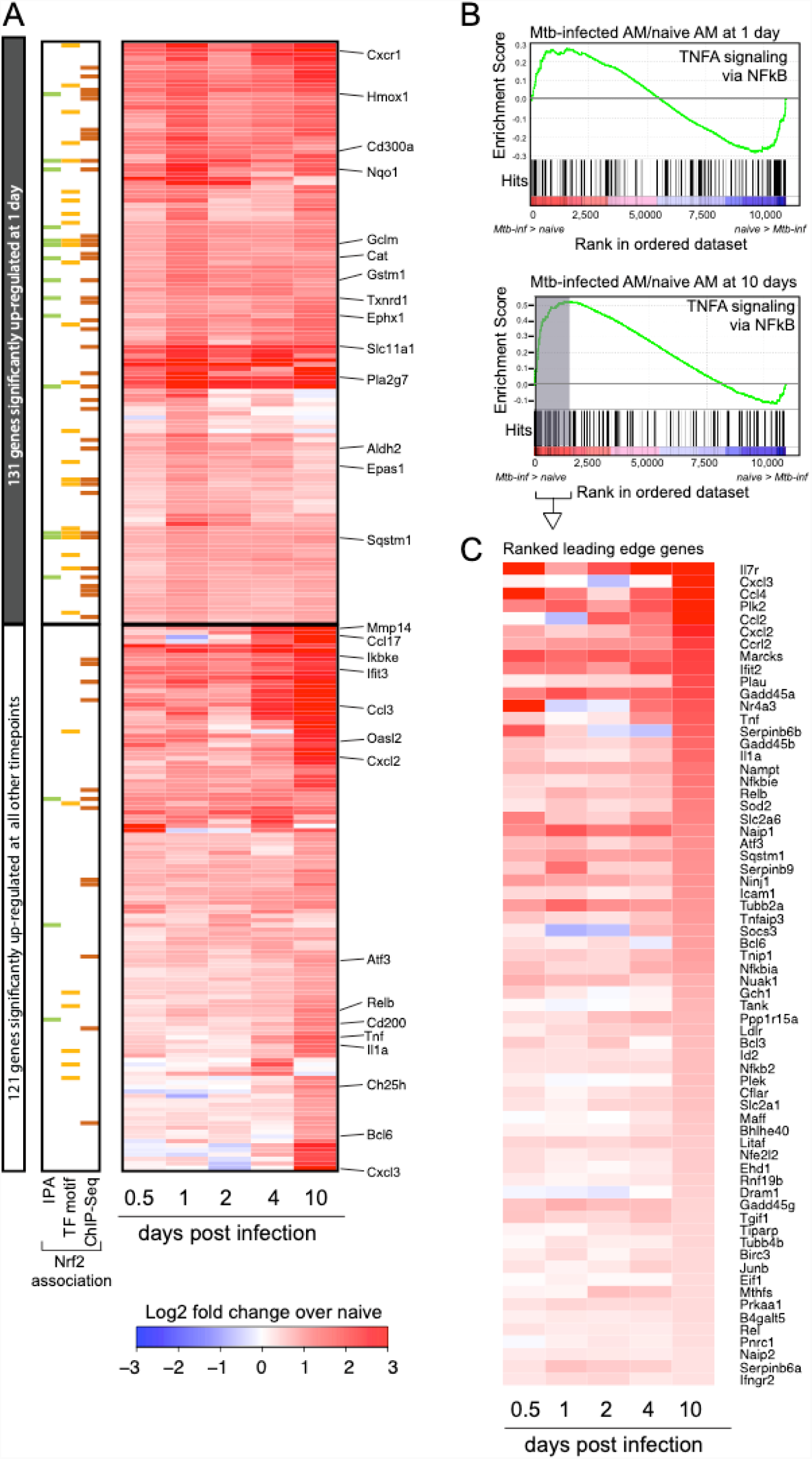
Over the first 10 days of infection, expression of Nrf2-associated genes is sustained and expression of pro-inflammatory genes is delayed in Mtb-infected alveolar macrophages. (A) Heatmap of gene expression (log_2_ fold change over naïve AMs) for 252 genes up-regulated in Mtb-infected AMs compared to naïve AMs for at least one out of five time points (Filtering criteria: average CPM >1, |fold change| > 2 and FDR < 0.01, Benjamini-Hochberg calculated). Top 131 genes are significantly up-regulated in Mtb-infected AMs at 1 day post-infection. Bottom 121 genes are not significantly up-regulated in Mtb-infected AMs at 1 day post-infection. Colored bars indicate Nrf2 association as described in *Figure 2*. (B, C) Gene set enrichment analysis and top 50 ranked leading edge genes in the “TNFA signaling via NFkB” pathway for Mtb-infected AMs at 10 days post-infection. Data is presented from 3 independent experiments per time point.

The kinetic analysis also identified a late pro-inflammatory response up-regulated in Mtb-infected AMs primarily at 10 days post-infection (**Fig 4A,** *bottom*). At 10 days post-infection, Mtb-infected AMs displayed significant up-regulation of genes in the TNFA signaling via NFkB Pathway as determined by Gene Set Enrichment Analysis (GSEA) compared to naïve AMs (NES = 1.8, FDR = 0.0013), a pathway not significantly enriched in Mtb-infected AMs 1 day following infection (**Fig 4B**). The ranked leading edge genes in this pathway reveal the changes in gene expression over time, including increases in expression of: *Il1a, Tnf, Rel, Relb, NFkb2, Ccl2, Bhlhe40,* and *Nfe2l2* (**Fig 4C**). A number of pro-inflammatory cytokine and chemokine genes (*Tnf, Il1a, Cxcl2, Cxcl3, Ccl17)* are significantly up-regulated only in Mtb-infected AMs 10 days after infection (**Fig S4**).

### Bystander AMs express a unique transcriptional signature 10 days after infection

To disentangle the responses to intracellular infection from the responses to the inflammatory milieu in the lung, we analyzed the transcriptomes of the bystander AMs (uninfected cells from infected animals) at 1 and 10 days post-infection. As described above, bystander AMs displayed no significant gene expression changes compared to naïve AMs at 1 day after infection (**Fig 5A**). In contrast, by 10 days after infection bystander AMs showed abundant changes in gene expression with a total of 205 significantly changed genes (**Fig 5B**). 28 of these genes (highlighted in blue) were shared with the Mtb-infected AMs, while 177 of them (highlighted in red) were uniquely changed in bystander but not in Mtb-infected AMs. Another 200 genes (highlighted in purple) were differentially expressed only in Mtb-infected AMs. Comparison between these sets of genes demonstrates that expression changes found only in Mtb-infected AMs are enriched for Nrf2-associated genes (**Fig 5D**), while expression changes found only in bystander AMs are enriched for other more inflammatory pathways (**Fig 5E**). These differences are confirmed by Ingenuity Pathway Analysis of the two datasets. While Nrf2-mediated oxidative stress response and PTEN signaling were more highly enriched in Mtb-infected AMs compared to bystander AMs, bystander AMs showed differential expression for genes in a number of other pathways including calcium signaling, NFAT regulation of the immune response, role of pattern recognition receptors in recognition of bacteria and viruses, and STAT3 pathway (**Fig 5F**). Enrichment of these pathways indicate that some of the inflammatory signals received by bystander cells appear to be blocked in infected cells 10 days after infection. Overall, these data suggest that over the first week and a half AMs respond directly to infection as well as to systemic changes in the lung environment and that these two signals may cross-regulate.

**Figure 5:**
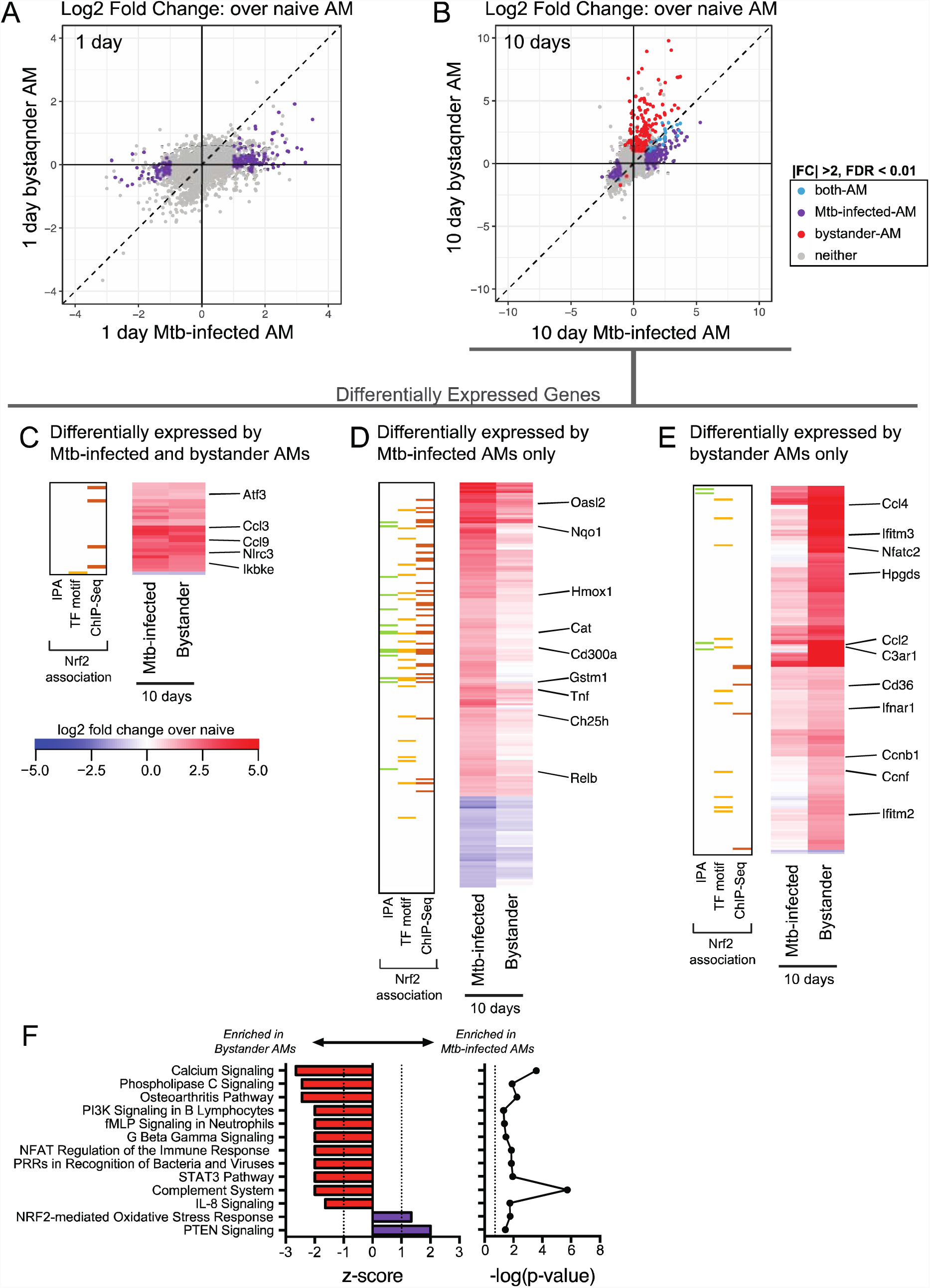
Bystander AMs express a unique transcriptional signature 10 days after infection. (A, B) Scatterplots comparing gene expression values (log_2_ fold change over average of naïve AMs) for H37Rv-infected versus bystander AMs at 1 day (A) and 10 days (B) post-infection with significant differentially expressed genes highlighted (|fold change| > 2 and FDR < 0.01, Benjamini-Hochberg calculated). Data is presented from 3 independent experiments. (C, D, E) Heatmaps of log2 fold change gene expression at 10 days over average of naïve AMs. Colored bars indicate Nrf2 association as described in *Figure 2*. (C) 28 genes differentially expressed by both bystander and Mtb-infected AMs. (D) 200 genes differentially expressed only by Mtb-infected AMs. (E) 177 genes differentially expressed only by bystander AMs. Columns represent the average of three independent experiments. Genes of interest noted to the right. (F) Ingenuity Pathway Analysis comparing gene expression from bystander AMs and Mtb-infected AMs. Canonical pathways with |z-scores| >1 and p-values < 0.05 were reported.

### Both cell-intrinsic and environmental factors shape the alveolar macrophage response to Mtb

To determine whether the AM response to Mtb is cell-intrinsic or environment-dependent, we compared the response of AMs infected *in vitro* to the *in vivo* measurements described above. We isolated AMs by BAL from naïve WT mice, allowed them to adhere for 18 hours, infected them with H37Rv, and measured their transcriptional response and ability to control bacterial growth. In parallel, we performed identical experiments with bone-marrow-derived macrophages (BMDMs), which have been used extensively, including by our group, to investigate how macrophages respond to Mtb (*35*). Similar to their response to Mtb infection *in vivo*, AMs displayed little to no increase in pro-inflammatory gene expression (including *Il1b, Il6* and *Nos2*) after infection *in vitro*, while BMDMs greatly up-regulated these genes (**Fig 6A**). One notable exception was *Tnf*, which was significantly up-regulated by AMs in response to H37Rv infection *in vitro*. In contrast to AMs infected *in vivo*, AMs infected *in vitro* did not up-regulated Nrf2-associated genes (**Fig 6B**). Overall, AMs were more permissive to bacterial growth than BMDMs, leading to a significant increase in bacterial burden as measured by CFU 5 days after infection (**Fig 6C**). These results suggest that the inability of AMs to up-regulate pro-inflammatory genes in response to intracellular infection is cell-intrinsic, while the up-regulation of the Nrf2-associated pathway is dependent on signals from the lung microenvironment.

**Figure 6:**
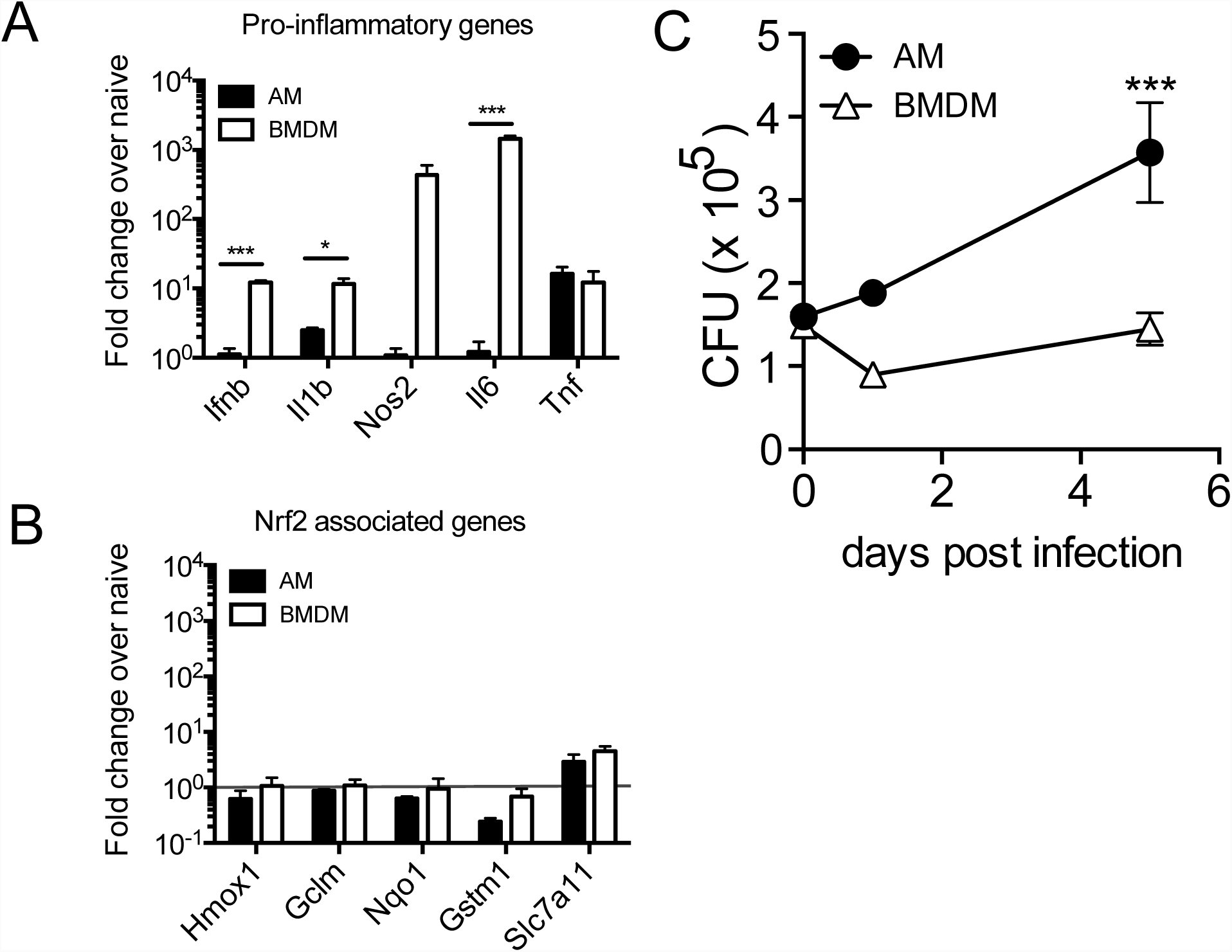
The alveolar macrophage response to Mtb is driven by both cell type and environment. *In vitro* H37Rv infection of AMs and bone marrow derived macrophages (BMDMs). (A) RT-qPCR gene expression analysis of pro-inflammatory genes 8 hours post-infection. (B) RT-qPCR analysis of Nrf2-associated genes 8 hours post-infection. (C) Colony forming unit (CFU) assay to measure bacterial burden in each cell type over 5 days. Data is representative of 3 independent experiments with three technical replicates each. Multiple t-tests with Holm-Sidak correction. *p<0.05, ***p < 0.001

### Expression of Nrf2 impairs the ability of AMs to control bacterial growth

Our computational analyses identified Nrf2 as a potential regulator of the *in vivo* AM response to Mtb-infection. To test this hypothesis, we isolated Mtb-infected AMs from Nrf2^-/-^ mice 24 hours after infection with mEmerald-H37Rv and performed RNA-sequencing. The response of Nrf2^-/-^ AMs was strongly attenuated compared to that of WT AMs and many of the genes exhibiting altered responses were associated with Nrf2 by IPA, transcription factor motif analysis, or ChIP-Seq analysis (**Fig 7A**). No additional genes to those identified in WT AMs were differentially expressed in Nrf2^-/-^ AMs (|fold change| > 2, FDR < 0.01) suggesting that Nrf2 does not act to restrain the transcriptional response in this setting, as has been reported previously (*36*).

**Figure 7:**
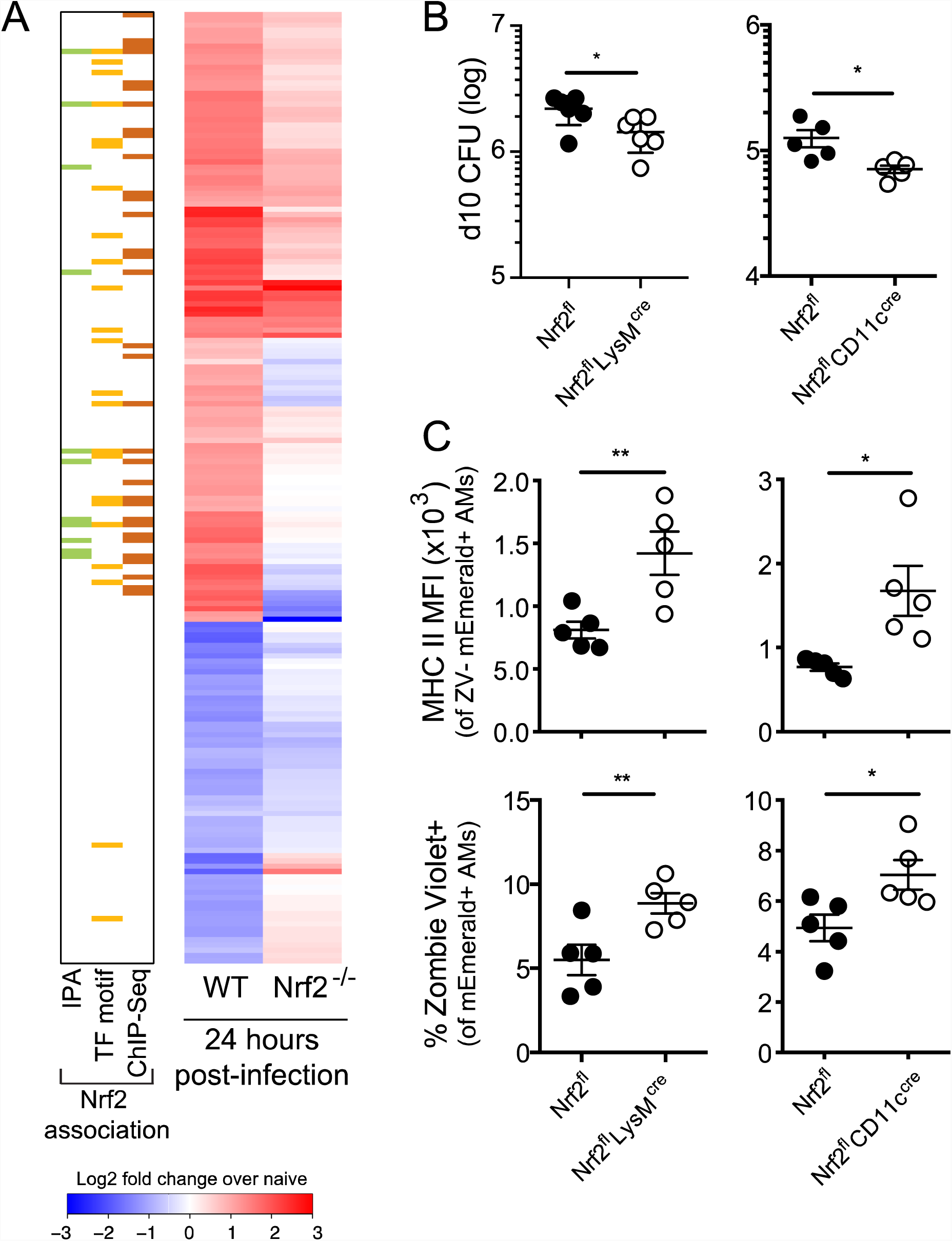
Modulation of Nrf2 activity alters macrophage response and control of Mtb. **(A)** Heatmap of gene expression (log_2_ fold change over average of respective naïve AMs) for WT and Nrf2^-/-^ Mtb-infected AMs, averaged from at least two independent experiments. Rows depict the 196 differentially expressed genes between naïve and Mtb-infected WT AMs as shown in *Figure 2B*. Colored bars indicate Nrf2 association as described in *Figure 2*. (B) Lung bacterial burden measured by CFU assay at 10 days post-infection with low dose H37Rv from Nrf2^fl^LysM^cre^, Nrf2^fl^CD11c^cre^ and their respective Nrf2^fl^ littermate controls. (C) MHC II MFI of live Mtb-infected AMs (*top*) and % dead (Zombie Violet Viability Dye^+^) of Mtb-infected AMs (*bottom*) as measured by flow cytometry at 10 days post-infection with high dose mEmerald-H37Rv from Nrf2^fl^LysM^cre^, Nrf2^fl^CD11c^cre^ and their respective Nrf2^fl^ littermate controls. Data is presented from 2 independent experiments (A) or representative of 2 independent experiments with 5 mice/group (B-C). Two-tailed unpaired Student’s t-test * p< 0.05, ** p < 0.01.

Nrf2 functions as a master regulator of an antioxidant stress response and likely plays critical roles in many cell types Our own flow cytometric analysis of *Nrf2*^*-/-*^ mice indicated that the cellularity and activation state of immune cells in their lungs are altered (data not shown); therefore, we chose to evaluate the functional role of Nrf2 in AMs during Mtb infection by generating Nrf2 conditional knockout mice. While there is no model for AM-specific gene deletion, we utilized both LysM^cre/+^ and CD11c^cre/+^ strains that delete floxed genes in either macrophages and neutrophils or AMs and dendritic cells, respectively (*37*), in combination with an Nrf2^floxed/floxed^ strain. These breedings generated the following conditional knockout models: Nrf2^floxed/floxed^; LysM^cre/+^ (Nrf2^fl^ LysM^cre^) and Nrf2^floxed/floxed^; CD11c^cre/+^ (Nrf2^fl^CD11c^cre^). We confirmed the absence of Nrf2 protein in AMs in both strains by Western Blot (**Fig S5**). We examined the bacterial burden in these mice at 10 days post-infection, the latest time point at which the vast majority of bacteria are still within AMs and found that both conditional knock-out models had lower bacterial burdens than their Nrf2^fl^ littermate controls (**Fig 7B).** At this time point, Mtb-infected AMs lacking Nrf2 expression were also more highly activated with up-regulation of MHC II surface expression and were more prone to cell death as measured by the viability dye Zombie Violet **(Fig 7C).** These results suggest that at least one mechanism by which Nrf2 impedes host control is by blocking the activation and cell death of Mtb-infected AMs, which might prevent bacteria from being taken up by more bactericidal cell types.

## Discussion

We find that in vivo infection of AMs with live virulent mycobacteria does not produce the kind of pro-inflammatory response previously observed in other macrophage infection models. Our data suggest that the initial host response to Mtb infection is relatively anti-inflammatory and may contribute to the extended time required for antigen transport to the draining lymph node and subsequent T cell activation and adaptive immune control (*2*). Previous studies, including our own, have described Mtb-infected macrophages *in vitro* up-regulating pro-inflammatory genes primarily driven by NF-κB or type I IFN signaling (*16, 18, 33, 35*). Here we describe a unique Nrf2-driven transcriptional signature expressed in Mtb-infected AMs *in vivo*. While Nrf2 and several of its target genes, notably *Hmox1*, have been shown to be up-regulated during chronic Mtb infection in response to cellular and oxidative stress (*38–40*), our results suggest that the Nrf2 signature in AMs is likely a general response to phagocytic activity rather than to bacterial infection. The lack of an Mtb-specific immune response is surprising given that AMs express almost all of the canonical pathogen-sensing machinery required to detect Mtb infection (*41*). In the context of a meta-analysis of 32 studies covering 77 different host-pathogen interactions, this transcriptional response falls far outside the norm (*42*).

The fact that the Nrf2 transcriptional response has not been reported previously, despite numerous published studies of Mtb-infected macrophages, highlights the importance of using *in vivo* systems to dissect lung-specific immunological events (*43*). Our results suggest that established *in vitro* models do not adequately replicate the initial host response to Mtb. In addition, it is worth noting that our observations that murine AMs become more pro-inflammatory over time in culture (data not shown) are in concordance with reports that the response of human AMs to Mtb infection evolves from a profile unique to AMs towards one that is more similar to that of monocytes the longer they are cultured *in vitro* (*44*).

Studies using Nrf2^-/-^ mice have demonstrated both protective (*S. pneumoniae, S. aureus*, RSV, and *P. aeruginosa*) and harmful (*H. influenzae*, Marburg virus) roles for Nrf2 during infection (*45–51*). To evaluate the functional role of Nrf2 in AMs during Mtb infection, we chose to use two conditional knockout models (LysM^cre^ and CD11c^cre^). Cre expression by these promoters also deletes Nrf2 in either neutrophils and MDMs (LysM) or dendritic cells (CD11c). However, during the first 10 days of infection following aerosol challenge Mtb almost exclusively infects AMs and therefore this approach interrogates the direct role of Nrf2 in Mtb-infected AMs early during infection. Conversely, beyond 10 days, significant numbers of neutrophils, MDMs and dendritic cells begin to participate in the response making it impossible to link the outcome of infection at later times directly to AMs. In fact, we observe divergent phenotypes in the Nrf2^fl^LysM^cre^ and Nrf2^fl^CD11c^cre^ strains at 14 and 28 days following infection (data not shown), which indicates unique roles for Nrf2 in other cell types participating in the immune response to Mtb-infection that will be the subject of future studies.

In addition to infection, lung injury caused by noncommunicable conditions such as chronic obstructive pulmonary disease (COPD), cigarette smoking, cystic fibrosis, and exposure to air pollution have all been shown to activate Nrf2 (*52, 53*). Therefore, the finding that Nrf2 expression hinders AMs from controlling intracellular pathogen growth provides one potential mechanism by which smoking and indoor air pollution contribute to TB risk (*54, 55*).

The finding that AMs do not mount an inflammatory and bactericidal response to Mtb infection is supported by transcriptional profiling we performed in collaboration with Peterson et al on Mtb residing within infected AMs *in vivo* using a novel bead-based method, Path-Seq, to enrich for bacterial transcripts (under revision, *Molecular Systems Biology*) (*56*). These studies show that infection of AMs *in vivo* fails to induce the robust Mtb stress response that is observed after infection of BMDMs *in vitro*. Unlike infection of BMDMs, which causes Mtb to dynamically regulate mycolic acid biosynthesis and increase expression of regulators such as DosR that respond to environmental stress, infection of AMs leads to very few transcriptional changes. Targeting AMs may be one of Mtb’s most important virulence strategies. In order to spread effectively via aerosol transmission of small numbers of bacteria, Mtb targets a macrophage subtype that is seemingly not activated by pathogen-sensing pathways and instead responds in a manner that safeguards against cellular damage. This unexpected cell-protective response provides one explanation for the puzzling observation, termed the “macrophage paradox” by Vance and Price, that intracellular pathogens often infect the very cell type bestowed with the effector functions to control pathogen growth (*57*). In addition, the observation that bystander AMs up-regulate several inflammatory pathways that are not expressed in Mtb-infected AMs from the same lung suggests that intracellular Mtb infection is impeding AMs from receiving or processing certain signals from the inflammatory environment. This suppression may be an active process by Mtb and warrants further study.

Given recent studies demonstrating that innate responses can be “trained” by prior exposure to stimuli such as BCG vaccination, influenza, or adenoviral infection (*58–60*), we wondered whether the initial AM response to Mtb could be similarly modified. Using a model of TB containment after intradermal infection that generates low-level inflammation in specific tissues, we are able to show that pre-exposure to low levels of cytokines allows AMs to mount a more robust and pro-inflammatory response to aerosol Mtb infection that is associated with enhanced protection (manuscript in preparation). These results indicate that the response of AMs can be beneficially modified by the inflammatory milieu. Traditional TB vaccine strategies generally aim to potentiate lung-homing Mtb-specific T cells during the first days of infection, yet our data suggest that to be effective during the first week, T cells would have to target infected macrophages that are essentially “hidden” as a consequence of up-regulating antioxidant rather than pro-inflammatory responses. We hypothesize that a dual platform that simultaneously generates T cell memory and “trains” AMs to be more inflammatory could be an effective vaccine strategy.

AMs express a number of inhibitory receptors at baseline that are thought to contribute to their immunoregulatory phenotype (*6*). We don’t yet understand how these other signaling pathways impact the AM response to Mtb or provide redundancy with the Nrf2 pathway. Our data suggest that Mtb-infected AMs undergo a shift towards an increasingly pro-inflammatory phenotype over the first 10 days of infection, but the underlying mechanisms and the subsequent events remain unknown. Is this change cell-intrinsic or dependent on uptake and recognition of bacteria by other cell types following AM cell death? Similarly, we don’t yet understand why a portion of the pro-inflammatory program expressed in bystander AMs at 10 days post-infection is absent in infected AMs. Are there additional inhibitory mechanisms apart from Nrf2 that specifically restrain Mtb-infected AMs? Finally, we don’t yet know how human AMs respond to Mtb upon initial infection *in vivo*. These studies were performed using mice living in specific-pathogen free facilities that are exposed to a very limited set of environmental factors, while humans are continuously exposed to a variety of airborne stimuli and pathogens that may alter the baseline state of AMs. How pre-exposure to different stimuli impacts human AM function, and therefore TB susceptibility, is not yet understood.

Here we show that AMs up-regulate an Nrf2-dependent cell protective transcriptional program in response to Mtb infection *in vivo* and fail to up-regulate canonical inflammatory pathways normally associated with intracellular infection and that this impairs the host’s control of bacterial growth. Further investigation into the intervening events between initial AM infection and the adaptive T cell response as well as how the AM response to Mtb can be modified by environmental exposure may facilitate a better understanding of the early events of TB infection and inform novel TB vaccine strategies.

## Materials and Methods

### Study Design

The aim of this study was to measure the immune response of AMs to intracellular Mtb infection *in vivo*. We characterized the transcriptional profile of murine Mtb-infected alveolar macrophages after aerosol infection by sorting cells and performing low-input RNA-sequencing. To determine the effect of the transcription factor Nrf2 on the response of AMs, we generated Nrf2 conditional knock-out mouse models (Nrf2^fl^LysM^cre^ and Nrf2^fl^Cd11c^cre^) and tested their responses against Nrf2^fl^ littermate controls. The number of replicates for each experiment is indicated in the figure caption. No animals were excluded in these studies.

### Mice

C57BL/6 and Nrf2^-/-^ (B6.129X1-*Nfe2l2*^*tm1Ywk*^/J) mice were purchased from Jackson Laboratories (Bar Harbor, ME). Nrf2^floxed^ (C57BL/6-*Nfe2l2*^*tm1.1Sred*^/SbisJ), CD11c^cre^ (B6.Cg-Tg(Itgax-cre)1-1Reiz/J) and LysM^cre^ (B6.129P2-*Lyz2*^*tm1(cre)Ifo*^/J) were purchased from Jackson Laboratories and bred to generate Nrf2^fl^CD11c^cre^ and Nrf2^fl^LysM^cre^ mice. Mice were housed and maintained in specific pathogen-free conditions at Seattle Children’s Research Institute and experiments were performed in compliance with the Institutional Animal Care and Use Committee. 6-12 week old male and female mice were used for all experiments, except for RNA-sequencing, which used only female mice for uniformity. Mice infected with Mtb were housed in a Biosafety Level 3 facility in an Animal Biohazard Containment Suite.

### *M. tuberculosis* Aerosol Infections and Lung Mononuclear Cell Isolation

Most aerosol infections were performed with a stock of wildtype H37Rv transformed with an mEmerald reporter pMV261 plasmid, generously provided by Dr. Chris Sassetti and Christina Baer (University of Massachusetts Medical School, Worcester, MA). Some infections used an Mtb strain with a deletion of the virulence determinant RD1 region (ΔRD1), provided by Dr. David Sherman (SCRI, Seattle, WA), and transformed with the same mEmerald expression plasmid. For both standard (∼100 CFU) and high dose (∼2,000-4,000 CFU) infections, mice were enclosed in an aerosol infection chamber (Glas-Col) and frozen stocks of bacteria were thawed and placed inside the associated nebulizer. To determine the infectious dose, three mice in each infection were sacrificed one day later and lung homogenates were plated onto 7H10 plates for CFU enumeration, as previously described (*61*).

### Bead aerosolization

Carboxylate 1.0 µm fluorescent beads (ThermoFisher) were aerosolized using a LC Sprint Resuable Nebulizer (PARI) attached to a vacuum pump and an air flow regulator as described previously (Schroeder WG, 2009, Biotechniques).

### Lung Single Cell Suspensions

At each time point, lungs were removed and single-cell suspensions of lung mononuclear cells were prepared by Liberase Blendzyme 3 (70 ug/ml, Roche) digestion containing DNaseI (30 µg/ml; Sigma-Aldrich) for 30 mins at 37°C and mechanical disruption using a gentleMACS dissociator (Miltenyi Biotec), followed by filtering through a 70 µM cell strainer. Cells were resuspended in FACS buffer (PBS, 1% FBS, and 0.1% NaN_3_) prior to staining for flow cytometry.

### Alveolar Macrophage Isolation

Bronchoalveolar lavage was performed by exposing the trachea of euthanized mice, puncturing the trachea with Vannas Micro Scissors (VWR) and injecting 1 mL PBS using a 20G-1” IV catheter (McKesson) connected to a 1 mL syringe. The PBS was flushed into the lung and then aspirated three times and the recovered fluid was placed in a 15mL tube on ice. The wash was repeated 3 additional times. Cells were filtered and spun down. For antibody staining, cells were suspended in FACS buffer. For cell culture, cells were plated at a density of 1 x 10^5^ cells/well (96-well plate) in complete RPMI (RPMI plus FBS (10%, VWR), L-glutamine (2mM, Invitrogen), and Penicillin-Streptomycin (100 U/ml; Invitrogen) and allowed to adhere overnight in a 37°C humidified incubator (5% CO_2_). Media with antibiotics were washed out prior to infection with *M. tuberculosis*.

### Cell Sorting and Flow Cytometry

Fc receptors were blocked with anti-CD16/32 (2.4G2, BD Pharmingen). Cell viability was assessed using Zombie Violet dye (Biolegend). Cells were suspended in 1X PBS (pH 7.4) containing 0.01% NaN_3_ and 1% fetal bovine serum (i.e., FACS buffer). Surface staining included antibodies specific for murine: Siglec F (E50-2440, BD Pharmingen), CD11b (M1/70), CD64 (X54-5/7.1), CD45 (104), CD3 (17A2, eBiosciences), CD19 (1D3, eBiosciences), CD11c (N418), I-A/I-E (M5/114.15.2), and Ly6G (1A8) (reagents from Biolegend unless otherwise noted). Cell sorting was performed on a FACS Aria (BD Biosciences). Sorted cells were collected in complete media, spun down, resuspended in Trizol, and frozen at −80°C overnight prior to RNA isolation. Samples for flow cytometry were fixed in 2% paraformaldehyde solution in PBS and analyzed using a LSRII flow cytometer (BD Biosciences) and FlowJo software (Tree Star, Inc.).

### RNA-sequencing and Analysis

RNA isolation was performed using TRIzol (Invitrogen), two sequential chloroform extractions, Glycoblue carrier (Thermo Fisher), isopropanol precipitation, and washes with 75% ethanol. RNA was quantified with the Bioanalyzer RNA 6000 Pico Kit (Agilent). Due to the low number of Mtb-infected cells recovered (∼2,000-4,000 cells total after pooling BAL from 10-12 mice), all cDNA libraries were constructed and amplified using the SMARTer Stranded Total RNA-Seq Kit (v1 or v2) - Pico Input Mammalian (Clontech) per the manufacturer’s instructions. Libraries were amplified and then sequenced on an Illumina NextSeq (2 x 75, paired-end). Stranded paired-end reads of length 76 were preprocessed: The first three nucleotides of R1 (v1 kit) or R2 (v2 kit) were removed as described in the SMARTer Stranded Total RNA-Seq Kit - Pico Input Mammalian User Manual (v1: 112215, v2: 063017) and read ends consisting of 50 or more of the same nucleotide were removed). The remaining read pairs were aligned to the mouse genome (mm10) + Mtb H37Rv genome using the gsnap aligner (v. 2016-08-24) allowing for novel splicing. Concordantly mapping read pairs (average 15-million / sample) that aligned uniquely were assigned to exons using the subRead program and gene definitions from Ensembl Mus_Musculus GRCm38.78 coding and non-coding genes. Only genes for which at least three samples had at least 10 counts and had an average CPM > 1.0 were retained, resulting in a total of 10,946 genes. Differential expression was calculated using the edgeR package from bioconductor.org. False discovery rate was computed with the Benjamini-Hochberg algorithm (**Tables S2, S3**). Hierarchical clusterings were performed in R using ‘TSclust’ and ‘hclust’ libraries. Heat map and scatterplot visualizations were generated in R using the ‘heatmap.2’ and ‘ggplot2’ libraries, respectively.

### Ingenuity Pathway Analysis (IPA)

IPA (QIAGEN) was used to identify enriched pathways for differentially expressed genes between naïve and Mtb-infected or naïve and bystander AMs (cut-off values: FDR < 0.01, |FC| > 2) at various timepoints following infection. Canonical pathways with enrichment score −log(p-value) ≥ 5.0 are reported. IPA was also used to identify differentially enriched pathways between bystander and Mtb-infected AMs at 10 days post-infection (cut-off values: FDR < 0.05, |FC| > 2). Canonical pathways with |z-scores| >1 and p-values < 0.05 were reported.

### Promoter Scanning (HOMER)

Promoter regions of genes that were up-regulated (|FC| > 2, FDR < 0.01, log_2_(average counts per million) > 1.0) were scanned for DNA protein-binding motif over-representation using the HOMER program (v4.9.1, homer.salk.edu) (*29*). Promoter regions were defined as 2000 nucleotides upstream of the gene start to 1000 nucleotides downstream. Background sequences were taken from the promoter regions of expressed genes defined by Ensembl Mus_Musculus GRCm38.78 (N=10,946). 402 known motifs were scanned and hypergeometric p-values computed.

### Nrf2 ChIP-seq Analysis

Fastq files from the GEO data set GSE75175 were downloaded for three Nrf2 ChIP-seq experiments assaying peritoneal macrophages from wild-type mice including the corresponding sequencing of input DNA (*31*). For each sample, single ended reads of length 101 were filtered to remove those consisting of 50 or more of the same nucleotide or of low-quality base calls. Filtered reads were aligned against mm10 using gsnap (v. 2011-11-20) with no allowance for splicing. Uniquely mapped reads were filtered for duplicates based on alignment position. A total of 3,975 peaks were called using MACS2 (v.2.1.0) for the combined ChiP-seq samples using the input DNA libraries as the controls. Called peaks were annotated by checking for overlap with promoter regions as described above.

### Gene Set Enrichment Analysis (GSEA)

Input data for GSEA consisted of lists, ranked by −log(p-value), comparing RNAseq expression measures of target samples and naïve controls including directionality of fold-change. Mouse orthologs of human Hallmark genes were defined using a list provided by Molecular Signatures Database (MSigDB) (*62*). GSEA software was used to calculate enrichment of ranked lists in each of the respective hallmark gene lists, as described previously (*63*). A nominal p-value for each ES is calculated based on the null distribution of 1,000 random permutations. To correct for multiple hypothesis testing, a normalized enrichment score (NES) is calculated that corrects the ES based on the null distribution. A false-discovery rate (FDR) is calculated for each NES. Leading edge subsets are defined as the genes in a particular gene set that are part of the ranked list at or before the running sum reaches its maximum value.

### BMDM Isolation and Culture

Bone marrow-derived macrophages (BMDMs) were cultured in complete RPMI with recombinant human CSF-1 (50 ng/ml; PeproTech Inc.) for 6 days. Media with antibiotics were washed out prior to infection with H37Rv and replaced with complete RPMI without antibiotics.

### Mtb *In Vitro* Culture and Infection

H37Rv was grown in 7H9 media at 37°C to O.D. of 0.1-0.3. The final concentration was calculated based on strain titer and bacteria was added to macrophages at an effective multiplicity of infection (MOI) of 0.5 for two hours. Cultures were then washed three times to remove extracellular bacteria. Infected macrophages were cultured for up to 7 days. For CFU measurement, cells were lysed with 1% Triton X-100/PBS and lysate from triplicate conditions were plated in serial dilutions on Middlebrook 7H10 agar plates (ThermoFisher Scientific) and cultured at 37°C for 21 days.

### qRT-PCR

For gene expression analysis, 1 x 10^5^ AMs or BMDMs were plated in 96-well plates overnight, followed by *in vitro* infection as described above. RNA was isolated from cells using TRIzol (Invitrogen), two sequential chloroform extractions were performed, Glycoblue (10 μg; ThermoFisher) was added as a carrier, and RNA was precipitated with isopropanol and then washed with 75% ethanol. Quantitative PCR reactions were carried out using TaqMan primer probes (ABI) and TaqMan Fast Universal PCR Master Mix (ThermoFisher) in a CFX384 Touch Real-Time PCR Detection System (BioRad). Data were normalized by the level of EF1a expression in individual samples. Fold induction was computed with respect to the normalized expression levels of respective macrophages under unstimulated conditions within the same experiment.

### Microscopy

Cells acquired by BAL were spun down and resuspended in complete RPMI without antibiotics and plated on a Lab-Tek II 8-chamber glass-bottom slide (Nunc). The slides were kept for 2 hours in a 37°C humidified incubator (5% CO_2_) to allow adherence before the media was removed and the cells were fixed with 2% paraformaldehyde in PBS. The slides were blocked with 5% FBS in PBS for 1 hour at room temperature before surface staining with APC conjugated anti-CD45 (clone 30-F11; Biolegend) for 1 hour at room temperature. The slides were washed with PBS + 5% FBS for 5 minutes at room temperature for a total of three times before mounting with ProLong Diamond Antifade Mountant with DAPI (Thermo Fisher). Cells were imaged using a 100X objective (1.40 NA) on a DeltaVision Elite with the following excitation filter cubes: Cy5 (632/22), GFP (475/28), and DAPI (390/18) and emission cubes: Cy5(679/34), GFP (525/48), and DAPI (435/48). The entire cell volume was captured using a series of Z-stack images with a 0.2um step size. Number of *Mycobacteria* cells per macrophage were enumerated by manual counting using the Z-stack of images. Representative images were first deconvolved with a theoretical point spread function using SVI Huygens Essential before a maximum intensity project image was created in Imaris Image Analysis Software (Bitplane).

### Western Blot

Naïve AMs were isolated by BAL and pooled from 5 mice, plated overnight for adherence and stimulated for 5 hours with the Nrf2 agonist, dimethyl fumarate (3.65 μM; Sigma-Aldrich) prior to protein collection with RIPA buffer (Cell Signaling Technology) and HALT protease inhibitor (ThermoFisher). Western blotting analyses were performed using standard techniques and transblotted onto nitrocellulose membranes. Membranes were probed with relevant primary antibodies: rabbit anti-Nrf2 (D1Z9C) (1:1000; Cell Signaling Technology), rabbit anti-mouse beta-actin1-HRP antibody (1:5000, Jackson Immunoresearch). Primary antibodies were detected by a secondary rat anti-rabbit-HRP antibody (1:2000, Jackson Immunoresearch).

### Statistical Analyses

RNA-sequencing was analyzed using the edgeR package from Bioconductor.org and the false discovery rate was computed using the Benjamini-Hochberg algorithm. All other data are presented as mean ± SEM and analyzed by one-way ANOVA (95% confidence interval) with Dunnett’s post-test (for comparison of multiple conditions) or unpaired Student’s t-test (for comparison of two conditions). Statistical analysis and graphical representation of data was performed using either GraphPad Prism v6.0 software or R. At least 3-5 mice were used per group in each experiment and all experiments were performed at least 2-3 times, as indicated in the figure legends. For p-values, * p < 0.05, ** p < 0.01, *** p < 0.001.

## Acknowledgements

We thank the staff at Seattle Children’s Research Institute vivarium for animal care, Pamela Troisch and the Next Gen Sequencing core at the Institute for Systems Biology, and Chris Sassetti and Christina Baer at the University of Massachusetts Medical School for the mEmerald bacterial strains. Members of the Urdahl lab provided helpful discussions.

## Author contributions

A.C.R, G.S.O., J.N., A.H.D., and A.A. designed the experiments. A.C.R., D.M., G.S.O, and J.N. conducted the experiments. L.M.A. and A.H.D. performed computational analyses. A.C.R., A.H.D., E.S.G., and A.A. wrote the paper.

## Data and materials availability

Raw and processed RNA-sequencing data can be accessed from the National Center for Biotechnology Information (NCBI) Gene Expression Omnibus (GEO) database under accession number [GSE: holder until GEO submission]. [*Submission pending.*]

## Supplementary Materials

Fig. S1. Lung flow cytometry gating scheme for detection of Mtb infected cells

Fig. S2. Gene expression heatmaps of naïve, bystander and Mtb-infected alveolar macrophages 24 hours post-infection

Fig. S3. Gating strategy for sorting of ΔRD1-infected AMs and bead-positive AMs

Fig. S4. Gene expression heatmaps of Mtb-infected alveolar macrophages 0.5, 1, 2, 4 and 10 days after aerosol infection

Fig. S5. Nrf2 Western Blot for alveolar macrophages from Nrf2^fl^LysM^cre^, Nrf2^fl^ CD11c^cre^, Nrf2^fl^ littermate controls, WT and Nrf2^-/-^ mice

Table S1. Ingenuity Pathway Analysis and HOMER results tables

Table S2: RNA-Sequencing data for WT AMs

Table S3: RNA-Sequencing data for Nrf2^-/-^ AMs

