## Supplemental Figures 1-5 for "Alveolar macrophages up-regulate a non-classical innate response to *Mycobacterium tuberculosis* infection *in vivo*"

**Supplementary Materials**

Fig. S1. Lung flow cytometry gating scheme for detection of Mtb infected cells

Fig. S2. Gene expression heatmaps of naïve, bystander and Mtb-infected alveolar macrophages 24 hours post-infection

Fig. S3. Gating strategy for sorting of ΔRD1-infected AMs and bead-positive AMs

Fig. S4. Gene expression heatmaps of Mtb-infected alveolar macrophages 0.5, 1, 2, 4 and 10 days after aerosol infection

Fig. S5. Nrf2 Western Blot for alveolar macrophages from Nrf2^fl^LysM^cre^, Nrf2^fl^ CD11c^cre^, Nrf2^fl^ littermate controls, WT and Nrf2^-/-^ mice

Table S1. Ingenuity Pathway Analysis and HOMER results tables

Table S2: RNA-Sequencing data for WT AMs

Table S3: RNA-Sequencing data for Nrf2^-/-^ AMs

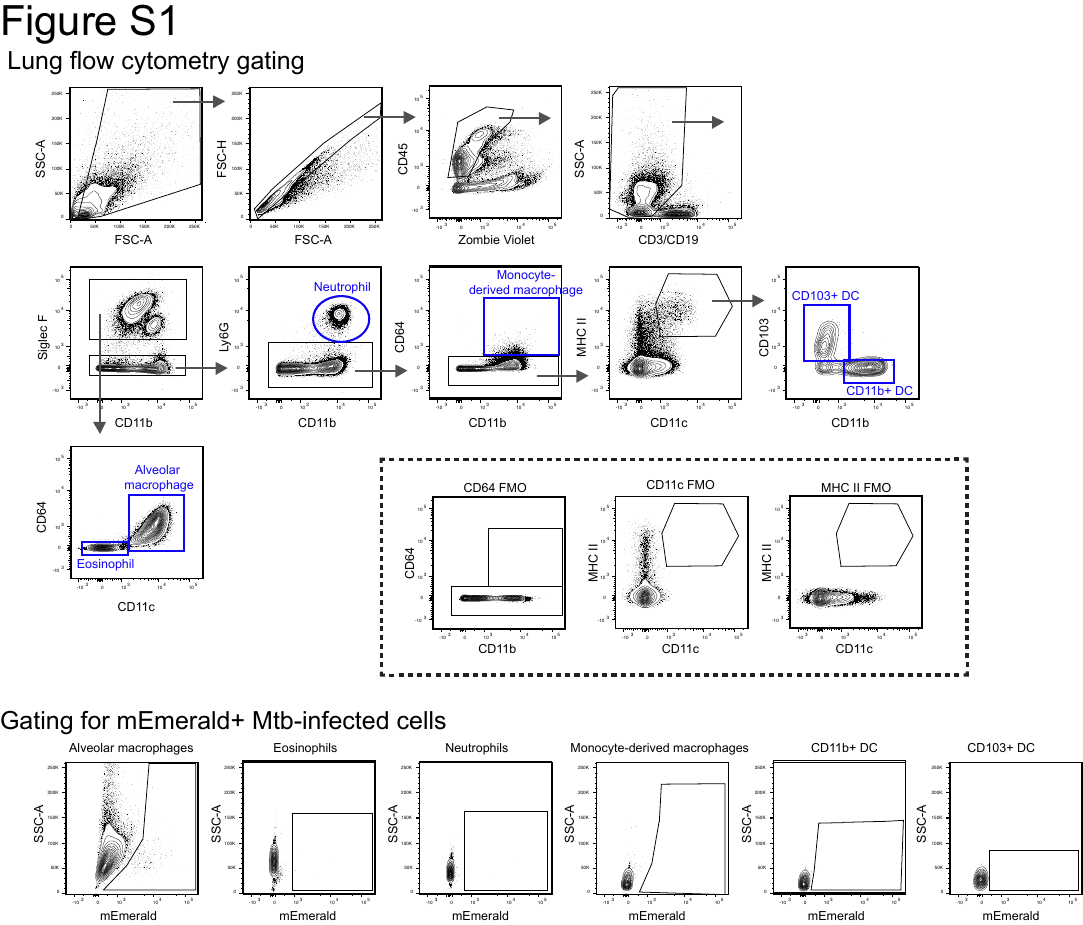

**Figure S1:** **Lung flow cytometry gating scheme for detection of Mtb infected cells.** *(Top)* Flow cytometry panel to detect alveolar macrophages, eosinophils, neutrophils, monocyte-derived macrophages, CD103^+^ and CD11c^+^ dendritic cells. Includes fluorescence minus one (FMO) controls for CD64, CD11c and MHC II staining. *(Bottom)* Gating for mEmerald^+^ Mtb-infected cells.

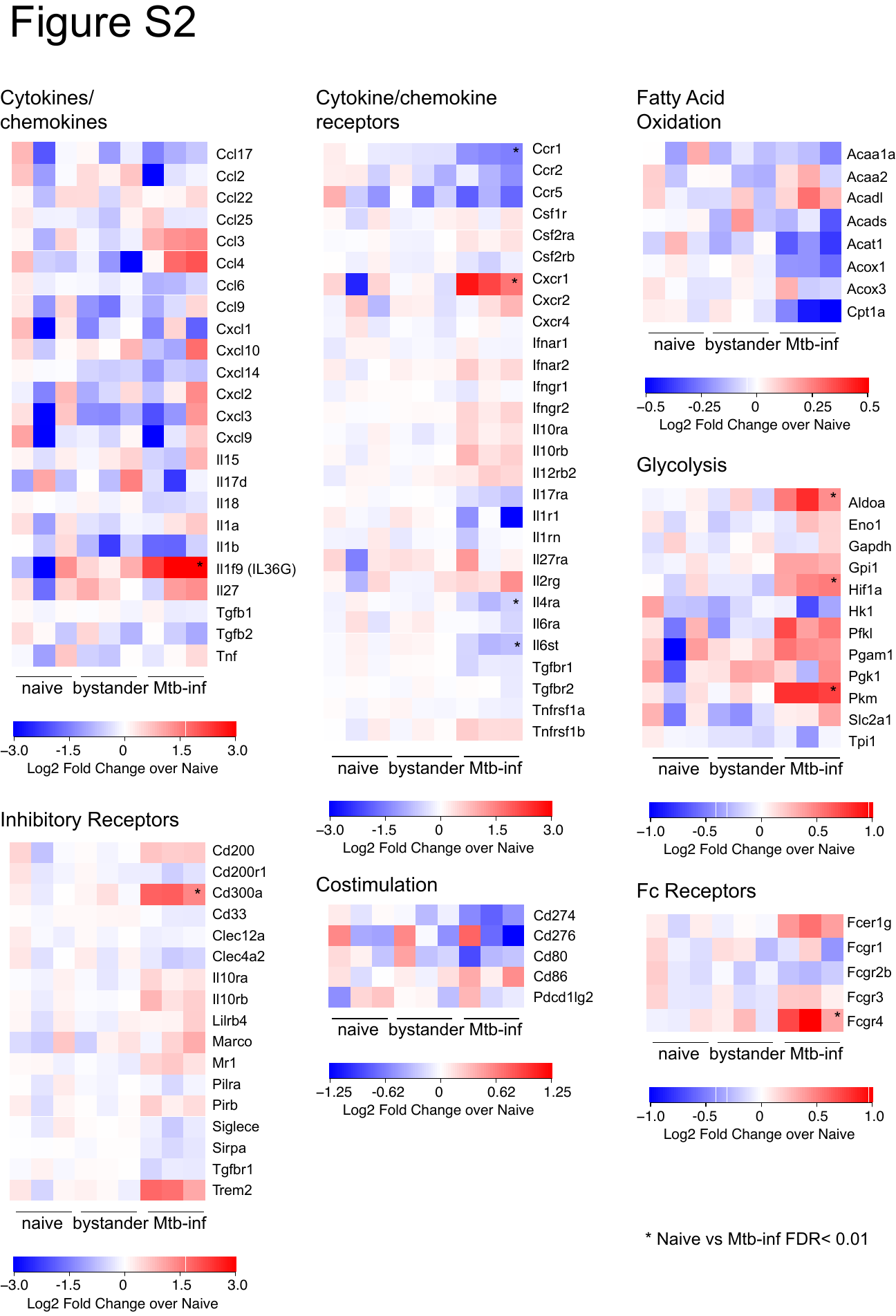

**Figure S2: Gene expression heatmaps of naïve, bystander and Mtb-infected alveolar macrophages 24 hours post-infection**. Log_2_ fold change gene expression over average of naïve AMs for relevant classes of genes. Data shown from 3 independent experiments. Asterisks indicate FDR < 0.01 (Benjamini-Hochberg calculated) for naïve versus Mtb-infected comparison.

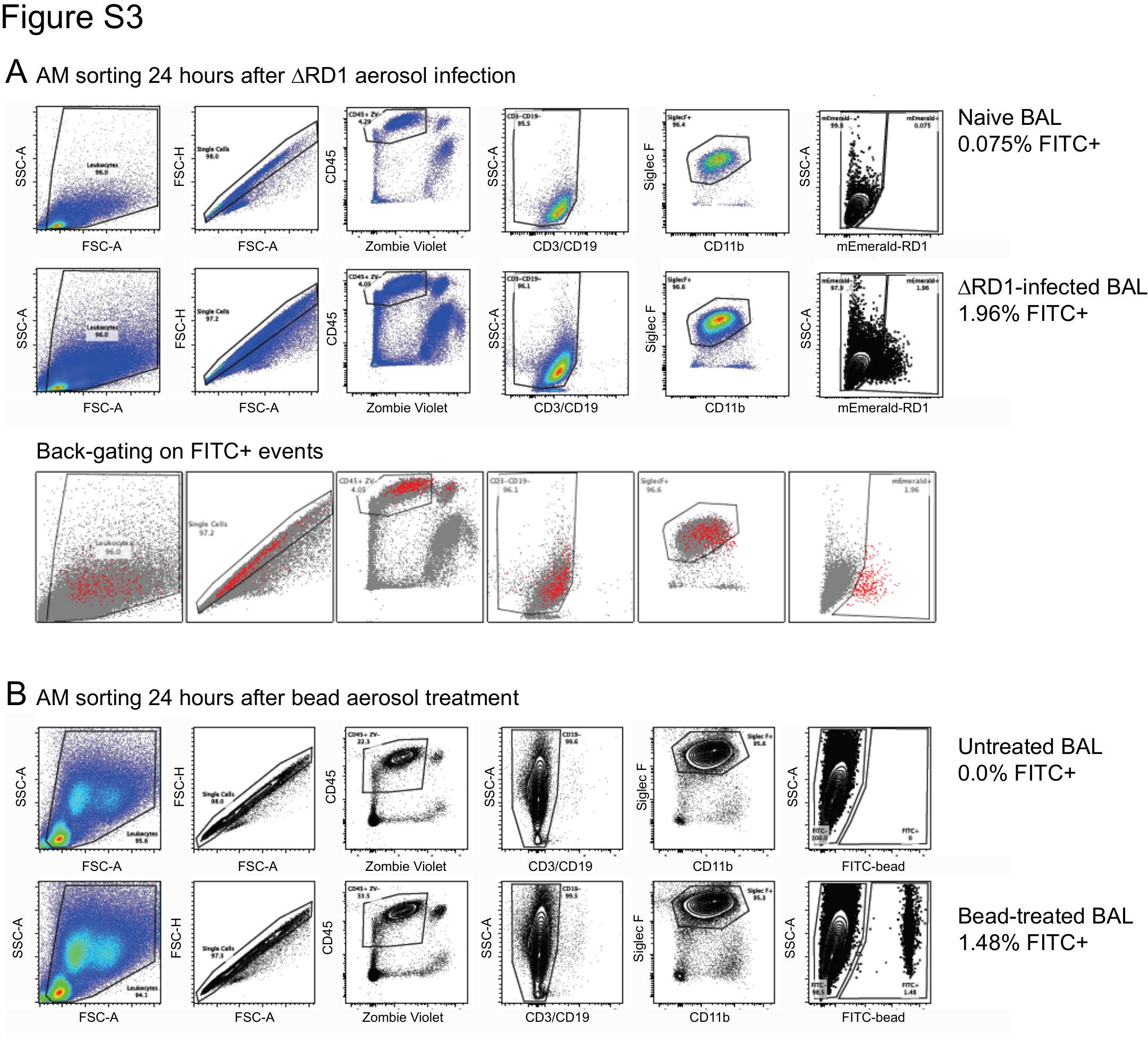

**Figure S3: Gating strategy for sorting of ΔRD1-infected AMs and bead-positive AMs.** (A) FACS sorting scheme for naïve and ΔRD1-infected AMs 24 hours after aerosol infection. Red dots indicate back-gating on FITC^+^ events. (B) FACS sorting scheme for naïve and bead-positive AMs 24 hours after bead aerosol treatment. Data is representative of 3 independent experiments.

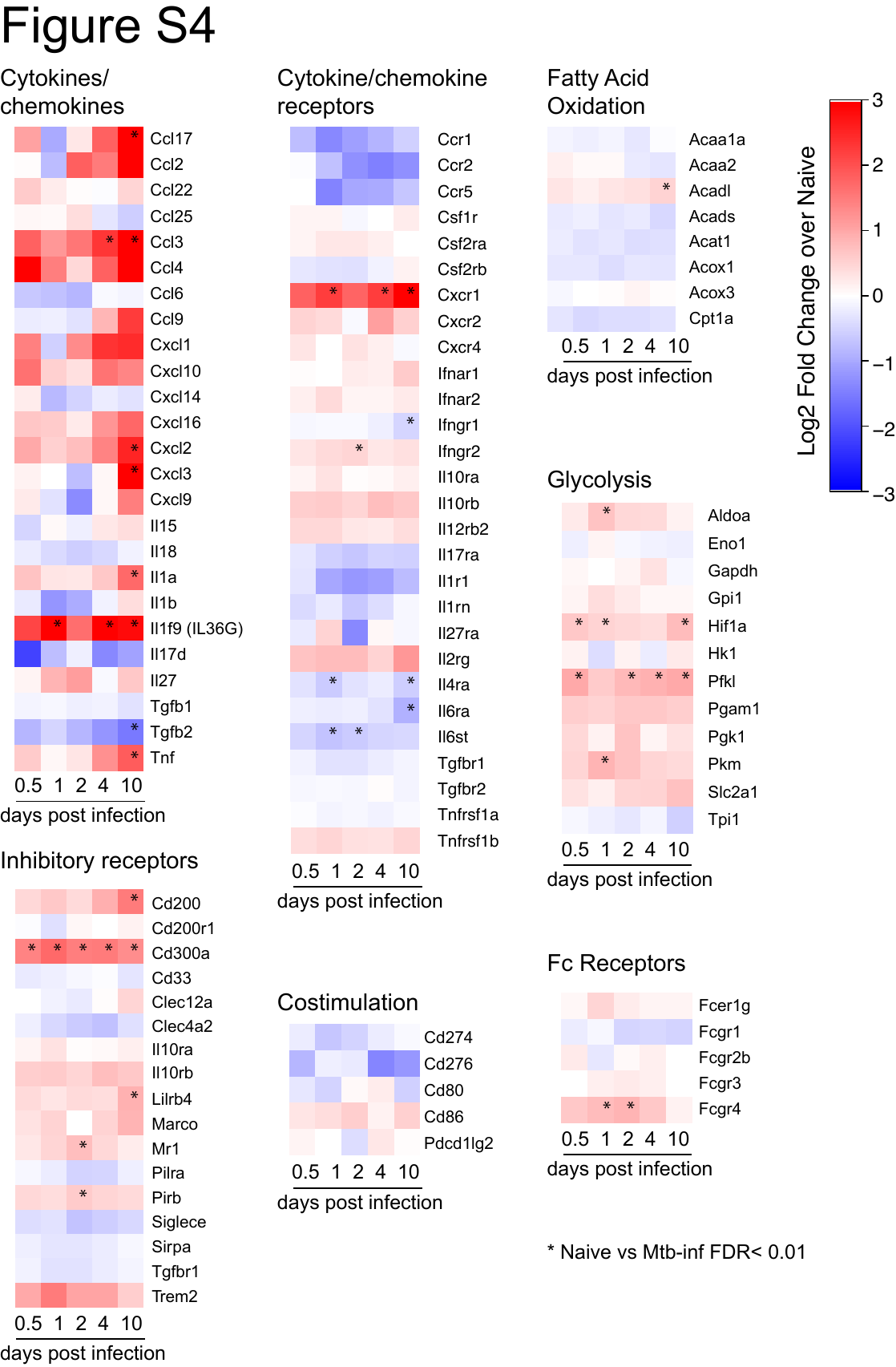

**Figure S4: Gene expression heatmaps of Mtb-infected alveolar macrophages 0.5, 1, 2, 4 and 10 days after aerosol infection.** Log_2_ fold change gene expression over average of naïve AMs for relevant classes of genes. Data shown is averaged from 3 independent experiments for each time point. Asterisks indicate FDR < 0.01 (Benjamini-Hochberg calculated) for naïve versus Mtb-infected comparison.

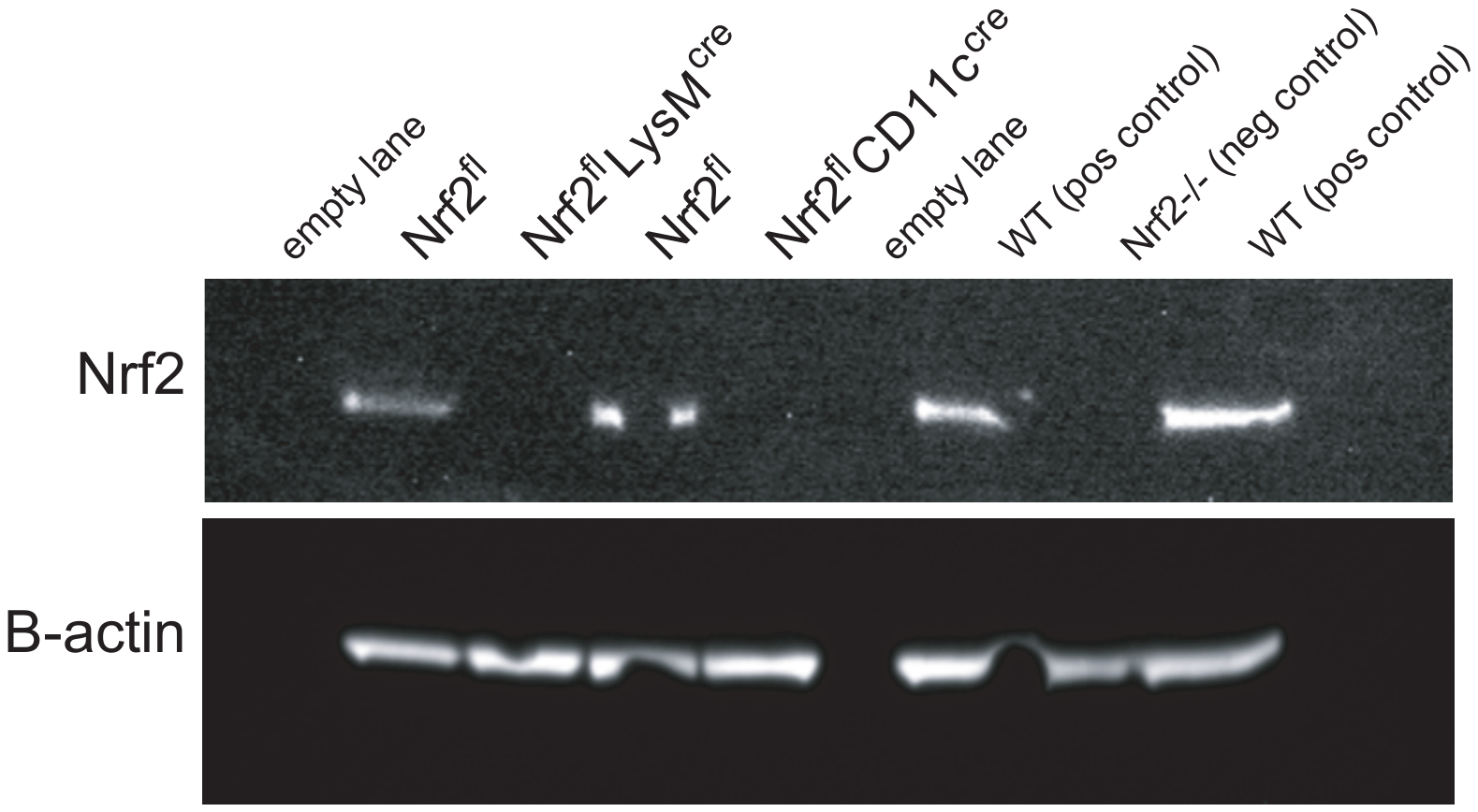

**Figure S5: Nrf2 Western Blot for alveolar macrophages from Nrf2^fl^LysM^cre^, Nrf2^fl^ CD11c^cre^, Nrf2^fl^ littermate controls, WT and Nrf2^-/-^ mice.**

Western Blots for Nrf2 and β-actin for AMs collected from indicated strains. Naïve AMs were isolated by BAL and pooled from 5 mice, plated overnight for adherence and stimulated for 5 hours with 3.65 μM dimethyl fumarate (Nrf2 agonist) prior to protein collection.

**Table S1: Ingenuity Pathway Analysis and HOMER results**

**Table S2: RNA-Sequencing data for WT AMs.** Normalized counts per million reads and fit coefficients for all expressed genes in WT AMs from H37Rv-infection 0.5-10 day time course, ΔRD1-infection at 24 hours, and bead-infection at 24 hours

**Table S3: RNA-Sequencing data for Nrf2^-/-^ AMs.** Normalized counts per million reads and fit coefficients for all expressed genes in Nrf2^-/-^ AMs from H37Rv-infection at 24 hours and WT AM controls.
